# Cytosolic linear DNA plasmids in *Saccharomycopsis* species

**DOI:** 10.1101/2023.07.20.549855

**Authors:** Eoin Ó Cinnéide, Padraic G. Heneghan, Arun S. Rajkumar, John P. Morrissey, Kenneth H. Wolfe

## Abstract

Some budding yeast species contain cytosolic linear DNA plasmids (also called virus-like elements, VLEs) that code for killer toxins that can kill other yeasts. The toxins are anticodon nucleases that cleave a specific tRNA in the cells being attacked, stopping translation. The best known plasmids of this type are the pGKL1/pGKL2 system of *Kluyveromyces lactis*. pGKL1 is a killer plasmid encoding the toxin zymocin (γ-toxin) which cleaves tRNA-Glu, and pGKL2 is a helper plasmid required for replication and transcription of pGKL1. Here, we investigated similar plasmids in the genus *Saccharomycopsis* that were originally described in the 1980s. *Saccharomycopsis* has undergone an evolutionary change of its genetic code, from CUG-Leu to CUG-Ser translation, which we hypothesized could have been driven by a tRNA-cleaving toxin encoded by a cytosolic plasmid. We sequenced a three-plasmid system in *S. crataegensis*, consisting of apparent killer, immunity, and helper plasmids. The killer plasmid contains genes coding for putative α/β (chitin-binding) and γ (ribonuclease) toxin subunits, but the γ-toxin gene is damaged in all the isolates we examined. We inferred the sequence of the intact *S. crataegensis* γ-toxin and expressed it in *Saccharomyces cerevisiae* and *Kluyveromyces marxianus*, but it did not cause a growth defect. We also identified free plasmids, or plasmids integrated into the nuclear genome, in nine other *Saccharomycopsis* species, including a case of recent interspecies transfer of a plasmid. Our results show that many yeasts in the CUG-Ser2 clade contain, or have in the past contained, plasmids related to those that carry anticodon nucleases.

## Introduction

As well as their nuclear and mitochondrial genomes, budding yeasts can contain other types of DNA and RNA genomes in their cytosol. These are usually called plasmids, viruses, or virus-like elements (VLEs) (Satwika et al., 2012a; Klassen et al., 2017; Schaffrath et al., 2018; Mannazzu et al., 2019).

There are three main types of these entities: circular double stranded DNA molecules (e.g. the 2- micron plasmid of *S. cerevisiae*), linear dsDNA molecules (e.g. the pGKL1 killer plasmid of *Kluyveromyces lactis*), and linear dsRNA molecules (e.g. the M1 killer plasmid of *S. cerevisiae*). The latter two encode killer toxins, first discovered for their killer phenotype, although the mechanisms of action of pGKL1 and M1 are different and they are inferred to have separate evolutionary origins. In this paper we focus on linear dsDNA plasmids because they are hypothesized to have been involved in evolutionary changes of the nuclear genetic code, as explained later in this section.

The *K. lactis* plasmids pGKL1 and pGKL2 comprise the best studied and understood cytosolic plasmid system (Gunge et al., 1981; Stark et al., 1990; Satwika et al., 2012a; Klassen et al., 2017; Schaffrath et al., 2018). pGKL1 (8.9 kb) is called a killer plasmid because it codes for the toxin. Zymocin is the name given to the complete toxin (holotoxin), which is a complex of three protein subunits called α, β, and γ. The γ subunit is the active subunit of the toxin, coding for the ribonuclease that cleaves the anticodon loop of tRNA-Glu(UUC), and is encoded by one of the four genes in pGKL1 (Hishinuma et al., 1984; Stark et al., 1984; Sor and Fukuhara, 1985; Lu et al., 2005). A second gene codes for the α and β subunits, which are synthesized as a single precursor protein that is later cleaved. Both the γ and α/β subunits have secretion signals at their N-termini and are secreted out of pGKL1-containing cells. They are covalently linked to each other by a disulphide bond near their C-termini (Wemhoff et al., 2014). The α/β protein contains a chitin-binding domain (glycosyl hydrolase GH18 domain, related to chitinase) and its function is to attach to the cell wall of the recipient cell and to deliver the active γ-toxin into that cell (Butler et al., 1991; Klassen et al., 2017). A third gene in pGKL1 codes for an immunity factor, which is not secreted and which prevents the toxin from cleaving tRNA in the host cell (Tokunaga et al., 1987; Kast et al., 2015; Klassen et al., 2017). pGKL2 (13.5 kb) is called a helper plasmid because it provides essential housekeeping functions without which pGKL1 cannot survive. Because these plasmids are located in the cytosol, they do not use the normal nuclear machinery for DNA replication and transcription. pGKL2 contains 11 genes including a DNA polymerase, an RNA polymerase, and a telomere-binding protein that is covalently attached to the 5’ ends of the linear DNA molecules (Tommasino et al., 1988; Satwika et al., 2012a).

The major target of zymocin was discovered to be the anticodon loop of tRNA-Glu(UUC), i.e. a glutamate tRNA with anticodon UUC (Lu et al., 2005). Subsequently, the toxins of three similar killer systems encoded by cytosolic linear DNA plasmids of other yeast species have been characterized. Two of these toxins, PaT of *Millerozyma acaciae* and DrT of *Debaryomyces robertsiae*, are also anticodon nucleases but they target a different tRNA, tRNA-Gln(UUG) (Klassen et al., 2008; Chakravarty et al., 2014; Klassen et al., 2014). The fourth characterized toxin, PiT of *Babjeviella inositovora*, has been shown to induce fragmentation of ribosomal rRNA rather than tRNA, cleaving 25S and 18S rRNAs at several sites (Kast et al., 2014). Each of these toxins is encoded by a killer plasmid with γ and α/β subunit genes and an immunity gene, assisted by a helper plasmid (Satwika et al., 2012a; Klassen et al., 2017). In the case of PiT there is also a third plasmid that appears to encode immunity functions.

In the 1980s and 1990s several laboratories used gel electrophoresis to screen natural isolates of numerous yeast species for the presence of cytosolic DNA plasmids. Bolen and colleagues discovered a set of three plasmids in *Saccharomycopsis crataegensis* by this method (Shepherd et al., 1987; Bolen et al., 1992). They described the plasmids as a ‘cryptic’ system because no killer activity was associated with them, although it can often be difficult to characterize natural activity of toxins because it can depend on abiotic factors such as pH. In addition, Fukuhara (1995) found natural plasmids by gel electrophoresis in two other *Saccharomycopsis* species, *S. malanga* and *S. fibuligera* (called *Botryoascus cladosporoides* in his paper). Fukuhara’s study was an exhaustive screen of over 1800 isolates (600 species) from the CBS (Westerdijk Institute) collection by gel electrophoresis, which resulted in the discovery of about 20 new plasmid systems in various species. He found that plasmids were present in only 1.5% of the isolates he screened and that even if a yeast species is the host of a natural cytosolic linear DNA plasmid, the plasmid is typically present in only a minority of strains of that species (Fukuhara, 1995).

*Saccharomycopsis* yeasts use a non-standard nuclear genetic code (Krassowski et al., 2018; Muhlhausen et al., 2018; Junker et al., 2019). They translate the codon CUG as serine, whereas most other organisms translate CUG as leucine. *Saccharomycopsis* and its sister genus *Ascoidea* form a group called the CUG-Ser2 clade, which is phylogenetically separate from the well known CUG-Ser1 clade of *Candida* species (including *C. albicans*) that also translates CUG as serine. The reassignments of CUG to serine in the CUG-Ser1 and CUG-Ser2 clades were the result of two separate evolutionary events on different branches of the phylogenetic tree. A third clade of yeasts reassigned CUG codons to alanine (CUG-Ala clade). These three evolutionarily parallel and separate changes in the nuclear genetic code all occurred in yeasts, which is remarkable because they are the only ones in eukaryotes (Krassowski et al., 2018). No other eukaryotes have undergone a ‘sense-to-sense’ codon reassignment, i.e. a reassignment of a sense codon from one amino acid to another, in nuclear genes.

The fact that all the known sense-to-sense genetic code reassignments in eukaryotes occurred in budding yeasts coincides conspicuously well with the fact that budding yeasts are the only eukaryotes known to possess anticodon nuclease toxins. Each of the three genetic code changes involved abandoning translation of CUG as leucine, which led our laboratory to hypothesize that natural selection to abandon this particular translation might have been caused by a killer toxin with an anticodon nuclease that specifically cleaved tRNA-Leu(CAG), the tRNA that translates CUG as leucine (Krassowski et al., 2018). Continual exposure to such a toxin could have imposed strong selection pressure to evolve alternative tRNAs that can translate CUG and are not cleaved by the toxin, which led to the emergence of serine or alanine tRNAs with CAG anticodons (Krassowski et al., 2018). Unlike in the CUG-Ser1 and CUG-Ala clades, the genomes of most species in the CUG-Ser2 clade contain genes for both the new tRNA (tRNA-Ser(CAG)) and the ancestral tRNA-Leu(CAG) (Ó Cinnéide et al., 2023), even though the latter is not used for translation except in one species, *Ascoidea asiatica* (Muhlhausen et al., 2018). The presence of both types of tRNA gene in the same genome suggests that the genetic code transition in the CUG-Ser2 clade is more recent than in the CUG-Ser1 and CUG-Ala clades, and we speculated that *Saccharomycopsis* or *Ascoidea* species could possibly still harbor a killer plasmid that drove the genetic code change away from CUG-Leu translation. In this scenario, a putative killer toxin γ subunit could exist that cleaves the ancestral tRNA-Leu(CAG) of budding yeasts.

For this reason we were interested in the previous reports of cytosolic linear DNA plasmids in *Saccharomycopsis*, and we set out to characterize them by whole-genome sequencing or data-mining of public data. We sequenced the plasmids of *S. crataegensis*, *S. malanga* and *S. fibuligera* that were described decades ago, and we found new plasmids in *S. selenospora* and in an unnamed *Saccharomycopsis* species. Furthermore, we discovered numerous instances of plasmids, or parts of plasmids, integrated into nuclear genomes throughout the *Saccharomycopsis* genus, mostly in species not harboring extant plasmids, indicating that the entire genus has probably been exposed to these cytosolic elements throughout its evolutionary history. We found a candidate toxin γ- subunit gene in *S. crataegensis* and characterized it using recombinant constructs in both *Saccharomyces cerevisiae* and *Kluyveromyces marxianus*, but did not detect killing activity. Finally, we present phylogenetic evidence from DNA polymerase genes of discrete subfamilies of circulating dsDNA plasmids. Overall, our study greatly expands the currently known diversity of yeast cytosolic linear DNA plasmid systems.

## Results and Discussion

### Genome sequencing and data analysis

We previously sequenced and assembled the genomes of the type strains of most species of the *Saccharomycopsis* genus as part of an earlier study (Ó Cinnéide et al., 2023). For the current study, we obtained specific other strains of *Saccharomycopsis* that were reported to contain cytosolic linear DNA plasmids in the studies of Bolen et al. (1992) or Fukuhara (1995), or on the Westerdijk Fungal Biodiversity Institute (https://wi.knaw.nl) web pages about individual strains (Table S1). In addition, we assembled Illumina raw data from 93 strains of *S. fibuligera* sequenced in a recent study (Wang et al., 2021). We searched these assemblies, and assemblies of publicly available *Saccharomycopsis* genomes, to find contigs containing homologs of known genes from cytosolic linear DNA plasmids.

We examined all contigs that had sequence similarity (TBLASTN hits) to genes of plasmid origin. In whole genome sequence assemblies, contigs that are derived from *bona fide* free-living cytosolic plasmids are expected to have two characteristic features: contig coverage that is much higher than for nuclear contigs (because the plasmids are usually present in high copy number), and terminal inverted repeats (TIRs) formed by the final 100-200 bp at each end of the linear genome (Satwika et al., 2012a). However, plasmids can also become integrated into the nuclear genome, either as complete plasmids or in part, and such integrations are relatively common in yeast genomes (Frank and Wolfe, 2009). They usually decay into pseudogenes as they age, but genes in recent integrations can be fully intact. Furthermore, free-living plasmids have some properties that can pose difficulties to genome assemblers: they are extremely A+T-rich; the DNA at their ends can be resistant to sequencing because it is covalently attached to the telomere-binding protein; and despite their small size they can contain repeated regions with nearly identical sequences (in the two TIRs, and between different plasmids in the same host). As a result, a plasmid’s sequence can sometimes be broken into multiple contigs in a genome assembly. Therefore we examined all *Saccharomycopsis* contigs that contained plasmid-like genes, regardless of whether they were derived from free-living plasmids or integrated into the nuclear genome, or even if their cellular location was unclear.

### *Saccharomycopsis* plasmids and plasmid-like sequences

1. *S. crataegensis* three-plasmid system.

Three cytosolic linear DNA plasmids were discovered in *S. crataegenesis* strain NRRL Y-5902 by Bolen and colleagues (Shepherd et al., 1987; Bolen et al., 1992) and named p*Scr*l-1, p*Scr*l-2, and p*Scr*l-3, with estimated sizes of 13.7, 7.0, and 5.8 kb respectively. p*Scr*l-2 and p*Scr*l-3 were found to share a region of high sequence similarity by Southern blotting. Two other *S. crataegensis* strains from the NRRL collection (Y-5903 and Y-5904) were reported to contain p*Scr*l-1 and p*Scr*l-2 but not p*Scr*l-3, and a further two strains (Y-5910 and YB-192) had no plasmids. Double-stranded RNA plasmids were also detected in each strain except Y-5902 (Shepherd et al., 1987).

We sequenced total cellular DNA from nine strains of *S. crataegensis* and found all three previously reported plasmids in four strains (Fig. 1A,B; Table S1). We discovered that strain NRRL Y-5902, which was used by Bolen et al. (1992) as a reference strain for analysis of the plasmids, contains a version of p*Scr*l-3 with a 4-kb deletion that reduces its size to 5.8 kb whereas it is 9.9 kb in NRRL YB-502. We therefore used NRRL YB-502 as a reference strain for the three plasmids instead, and we retain the name p*Scr*l-3 for the 9.9 kb plasmid (Fig. 1A). Our results confirm most of the results of Bolen et al. regarding which strains contain which plasmids (Table S1), except that we found that NRRL Y-5904 has a version of p*Scr*l-3 with a 6-kb deletion, as well as having p*Scr*l-1 and p*Scr*l-2 as previously reported (Bolen et al., 1992). All three *S. crataegensis* plasmids appear to have been lost from the Westerdijk Institute stocks of two strains (CBS 6447 and CBS 6448) whose NRRL counterparts (Y- 5902 and Y-5904) retain them. We also found that a Portuguese isolate, PYCC 8280, contains only the largest plasmid p*Scr*l-1 (Table S1), consistent with it being a helper plasmid. Our sequence data shows that all three *S. crataegensis* plasmids have terminal inverted repeats (TIRs). The region of high sequence similarity between p*Scr*l-2 and p*Scr*l-3 previously reported (Shepherd et al., 1987; Bolen et al., 1992) is 2.6 kb long and identical. It contains most of their DNA polymerase (DNAP) genes, which code for proteins with 98% identity, and a small downstream ORF (Dorf; Fig. 1A). There is also a DNAP gene on p*Scr*l-3, but it has only 47% amino acid identity to the other two DNAPs.

**Figure 1.**
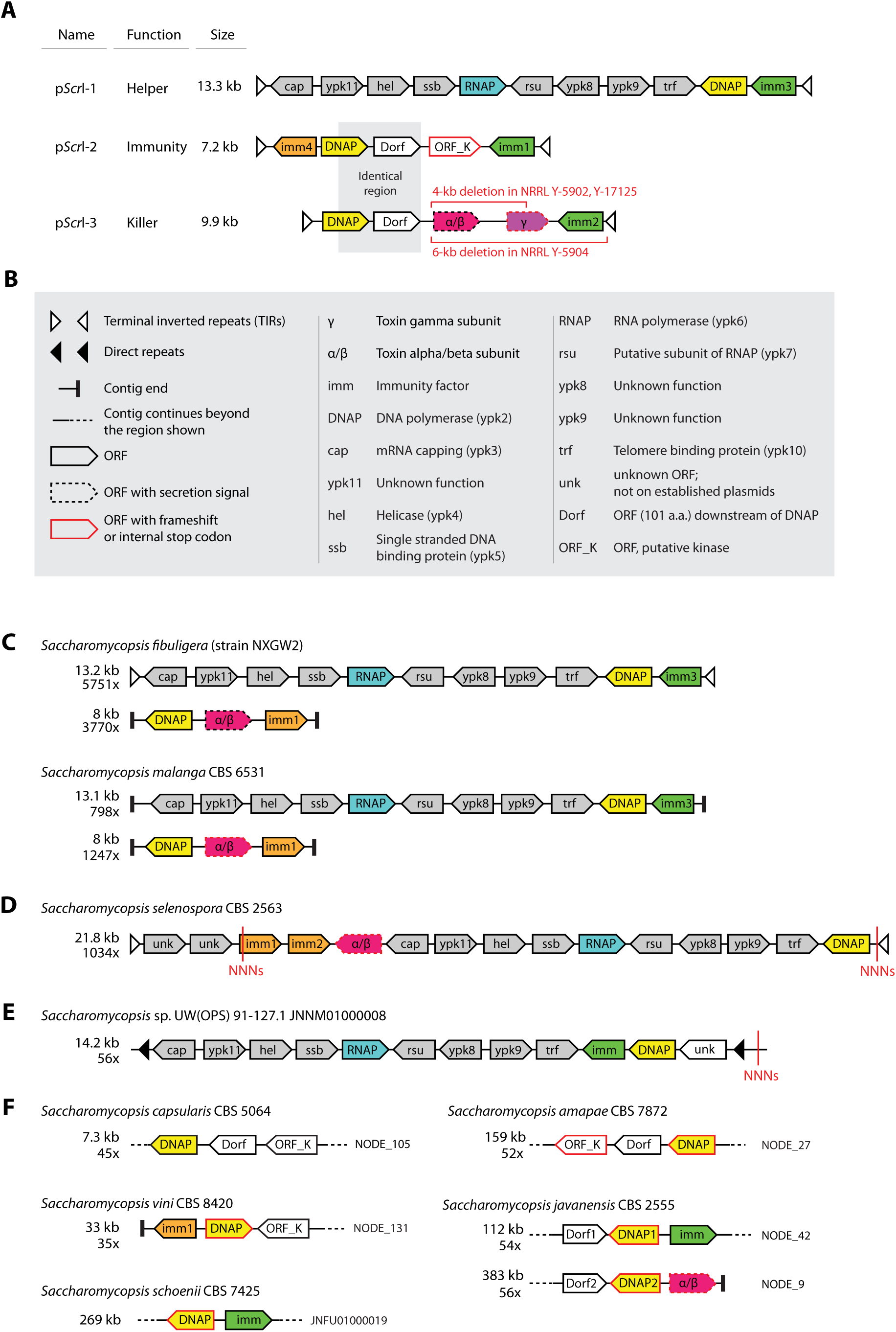
Maps of the *Saccharomycopsis* plasmids and plasmid-like contigs (not drawn to scale). (A) Genome organization of the three linear plasmids sequenced in *S. crataegensis*. Plasmid names (Bolen et al., 1992), their sizes in *S. crataegensis* strain NRRL YB-502, and their putative functions are shown. (B) Legend for plasmid maps and gene assignments, based largely on gene functions identified by experiments in *K. lactis* (summarized by Satwika et al. (2012a)). The *ypk* gene nomenclature follows Love et al. (2016). (C-F) Maps of newly discovered plasmid and plasmid-like sequences in other *Saccharomycopsis* species. Contig size and read coverage (where known) are shown. “NNNs” indicate short gaps in the sequence assembly, based on information from Illumina paired-end or mate-pair library sequencing.

The largest plasmid p*Scr*l-1 is inferred to be a helper plasmid with housekeeping functions similar to *K. lactis* pGKL2. It contains homologs of all 11 pGKL2 genes (Tommasino et al., 1988; Polomska et al., 2021), in almost the same order (Fig. 1A,B). The medium-sized plasmid p*Scr*l-3 (9.9 kb in strain YB- 502) is inferred to be a killer plasmid. It contains two genes with predicted secretion signals at their 5’ ends (predicted using TargetP (Emanuelsson et al., 2007)), coding for a toxin α/β (chitin-binding) subunit and a toxin ribonuclease γ-subunit. However the γ-toxin gene contains a frameshift mutation, discussed in more detail below. The 4-kb and 6-kb deletions in p*Scr*l-3 in three strains remove the whole α/β-subunit gene, and part or all of the γ-subunit gene (Fig. 1A; Table S1).

The smallest plasmid p*Scr*l-2 (7.2 kb) is predicted to have an immunity (antitoxin) function. We identified putative immunity genes in all three plasmids. These genes (*imm1*, *imm2* and *imm3*, colored green in Fig. 1A) have low but significant similarity to the immunity and immunity-like factors of *K. lactis* (pGKL1-ORF3 and pGKL2-ORF1), *D. robertsiae* (pWR1A-ORF5) and *M. acaciae* (pPac1-2-ORF4) (Kikuchi et al., 1985; Schaffrath et al., 1992; Chakravarty et al., 2014; Klassen et al., 2014; Kast et al., 2015). Each *S. crataegensis* plasmid contains one of these genes, with 31%-38% amino acid identity among them. In addition, p*Scr*l-2 contains a gene that we tentatively named *imm4*, which has significant amino acid sequence identity to plasmid genes of unknown function in *D. robertsiae* (pWR1A-ORF1) and *Schwanniomyces etchellsii* (pPE1A-ORF5) (Klassen et al., 2002; Klassen et al., 2004). We have detected numerous members of this gene family in other cytosolic linear plasmids, including in other *Saccharomycopsis* species as described below (colored orange in Fig. 1C-F). Although there is no experimental evidence that *imm4* and the other members of the ‘orange’ gene family encode an immunity function, its widespread distribution and the fact that some members of the ‘orange’ family have low but significant sequence similarity to members of the ‘green’ family (P.G.H., E. Ó C. and K.H.W., unpublished results), leads us to hypothesize that the ‘orange’ genes are also immunity genes.

We annotated two other genes in p*Scr*l-2. One (labeled ORF_K in Fig. 1A) codes for a 321 amino acid protein predicted to be a kinase, but it contains an internal stop codon. It has significant sequence similarity to predicted nuclear encoded kinases of *Ascoidea rubescens* (*Ascoidea* is the sister genus to *Saccharomycopsis*). In searches against the NCBI Conserved Domain Database (CDD ; Wang et al., 2023), ORF_K is identified as having a conserved phosphotransferase domain (Figure S1). This is the first instance of a kinase gene being found on such a cytosolic linear plasmid. The second gene (labeled Dorf in Fig. 1A) is a small ORF of 101 amino acids, located downstream of the DNAP gene, and it has an identical homolog in p*Scr*l-3. Database searches show that homologs of Dorf are present downstream of DNAP genes in cytosolic linear plasmids of several other species too, and we suspect that it may be the result of fragmentation of an original large DNA polymerase gene into two smaller genes (DNAP and Dorf) in some plasmids.

Horizontal transfer of a plasmid between two *Saccharomycopsis* species.

We identified two plasmids that are present at high copy number in several strains of *S. fibuligera* and one strain of *S. malanga* (Fig. 1C), consistent with the gel electrophoresis results of Fukuhara (1995). These are a 13-kb helper plasmid entirely collinear with *S. crataegensis* p*Scr*l-1, and an 8-kb plasmid containing three genes: a toxin α/β subunit gene, an immunity gene in the ‘orange’ family (related to *D. robertsiae* pWR1A-ORF1), and a DNAP gene (Fig. 1C). We also found that the entire 13- kb helper plasmid (including TIRs) has integrated into the nuclear genome of two strains of *S. fibuligera*: KJJ81 which is a hybrid strain (Choo et al., 2016), and strain ACX0001 which is not hybrid. The nuclear integration is present at the same place on chromosome 2 of ACX0001 and the ‘A’ copy of chromosome 2 of KJJ81, but absent from the ‘B’ copy, which suggests that the plasmid became integrated before the hybridization occurred.

Phylogenetic analysis of the 13-kb and 8-kb plasmid sequences from different strains (Fig. 2) leads to the conclusion that an interspecies transfer of an 8-kb plasmid must occurred between *S. fibuligera* and *S. malanga.* The 13-kb plasmid is present in three strains of *S. fibuligera* and these sequences differ by ≤ 5 nucleotides, whereas the 13-kb plasmid of *S. malanga* is 15% divergent from it.

**Figure 2.**
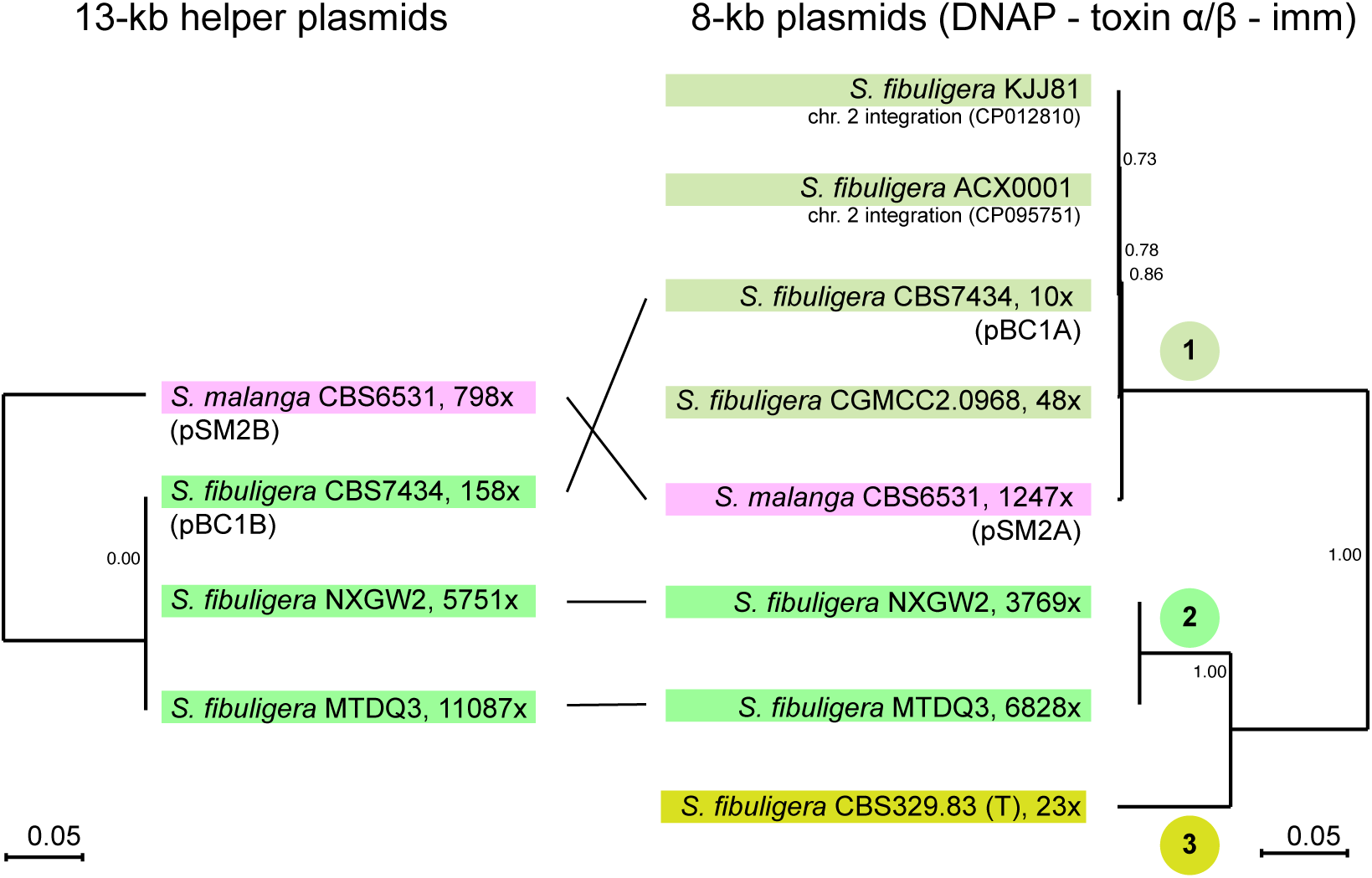
Comparison of phylogenetic trees constructed from the 13-kb and 8-kb plasmids found in strains of *S. fibuligera* and *S. malanga*. Lines in the centre indicate the 4 strains that contain both of the plasmids. Circled numbers 1-3 indicate the three divergent variants of the 8-kb plasmid present in *S. fibuligera* (scale bars indicate 5% DNA sequence divergence). For each plasmid, the coverage in our *de novo* assemblies is indicated by “x”. The plasmids are inferred to be free-living in all strains except *S. fibuligera* KJJ81 and ACX001 which have the 8-kb plasmid integrated into the nuclear genome (chromosome 2). The trees were constructed from DNA alignments of the complete plasmids, using MUSCLE and PhyML with their default settings in Seaview version 5.04 (Gouy et al., 2010). We confirmed that each genome assembly came from the species named, by BLASTN searches using ribosomal DNA queries. Plasmid names previously assigned by Fukuhara (1995) are indicated (pSM2A/B in *S. malanga* CBS6531, and pBC1A/B in *S. fibuligera* [*Botryoascus cladosporoides*] CBS7434).

However, there are three different versions of the 8-kb plasmid in *S. fibuligera*. They all have the same gene content but their sequences are significantly different from each other (10-15% divergence), forming three clades (Fig. 2), and the 8-kb plasmid in *S. malanga* has 99% DNA sequence identity to those in *S. fibuligera* clade 1. Within clade 1, there are approximately 50 nucleotide differences between *S. malanga* and each of the *S. fibuligera* strains, and ≤ 11 differences among the *S. fibuligera* sequences.

This situation must have arisen by interspecies transfer of an 8-kb plasmid, either (*i*) transfer of an 8- kb plasmid from *S. fibuligera* clade 1 into *S. malanga* CBS6531, or conversely (*ii*) transfer of an 8-kb plasmid from an *S. malanga* strain like CBS6531 into the common ancestor of *S. fibuligera* clade 1.

Comparing *S. malanga* CBS6531 to *S. fibuligera* CBS7434, their 8-kb plasmid are almost identical, but their 13-kb plasmids are 15% different in sequence. We also noticed that the toxin α/β subunit gene contains a frameshift in most of the 8-kb plasmids from these two species. It is a continuous ORF only in *S. fibuligera* CBS329.83 and NXGW2. It contains a single frameshift in each of the other six strains shown in the tree in Fig. 2, at three separate sites, so it may be non-functional in these six strains.

Plasmids and toxin α/β subunit genes in other *Saccharomycopsis* species.

In the *S. selenospora* assembly, we found a large (21.8 kb) plasmid with TIRs that is present at high copy number (Fig. 1D). This plasmid appears to be a fusion between a helper plasmid and a plasmid similar to the 8-kb plasmids of *S. fibuligera* and *S. malanga*. Its right end contains the 10 housekeeping genes normally found in helper plasmids, and its left end contains a predicted secreted toxin α/β subunit (with an internal stop codon), two predicted ‘orange’ family immunity genes related to *D. robertsiae* pWR1A-ORF1, and two ORFs of unknown function. We also found a complete helper plasmid integrated into the nuclear genome of the unnamed *Saccharomycopsis* species UWOPS 91-127.1 (Fig. 1E; Wendland and Hesselbart, NCBI accession number JNNM01000000).

As well as these essentially complete plasmids, we found lower-coverage contigs containing plasmid-like sequences in five other species: *S. capsularis*, *S. amapae*, *S. vini*, *S. javanensis* and *S. schoenii* (Fig. 1F). These contigs are inferred to come from the nuclear genome, based on their size and the presence of nuclear genes adjacent to the plasmid-derived genes. The *S. javanensis* contig includes a predicted toxin α/β subunit gene with an internal stop codon. These contigs were found primarily by TBLASTN searches using the *S. crataegensis* p*Scr*l-1 DNA polymerase protein as a query against all available *Saccharomycopsis* genomes. We detected many more nuclear integrants throughout the genus in these searches, as well as by TBLASTN searches using other large conserved plasmid proteins as queries, but many of the hits were heavily degraded pseudogenes and so are not shown in Fig. 1F. Such heavily degraded pseudogenes have been observed before across the subphylum Saccharomycotina (Frank and Wolfe, 2009; Satwika et al., 2012b). Interestingly, homologs of the putative p*Scr*1-2 kinase gene ORF_K are found adjacent to the DNA polymerase gene in several of these contigs (Fig. 1F).

### Reconstruction of *S. crataegensis* γ-toxin

Of all the plasmids we discovered in the *Saccharomycopsis* genus, only *S. crataegensis* p*Scr*l-3 contained a gene we inferred to be a likely γ subunit (ribonuclease) of a killer toxin (Fig. 1A).

However, the putative γ-toxin gene contains an apparent frameshift mutation in p*Scr*l-3 of *S. crataegensis* YB-502 that reduces its length and its similarity to known toxins, and the gene is partially deleted in all other strains that contain p*Scr*l-3 (Fig. 1A). By comparing the sequence of the frameshifted gene in *S. crataegensis* YB-502 to the incomplete γ-toxin gene in *S. crataegensis* Y- 17125, we identified a 2-nucleotide insertion in YB-502 as the probable cause of the frameshift and inferred a likely sequence of the full-length *S. crataegensis* γ-toxin protein, containing 268 amino acids (Fig. 3A,B). The inferred toxin protein sequence has all the canonical features of a *bona fide* anticodon nuclease toxin: an N-terminal secretion signal, a Glu residue near the N-terminus of the mature protein, and conserved His and Cys residues near the C-terminus (Fig. 3B). The protein sequences of the four currently known toxins are extremely divergent from one another, but these key residues are universally present and are known to be essential for toxin activity (Meineke et al., 2012; Chakravarty et al., 2014). The Glu and His residues are close to the active site of the ribonuclease (Chakravarty et al., 2014), while the Cys is required for cross-linking to the α/β subunit (Wemhoff et al., 2014).

**Figure 3.**
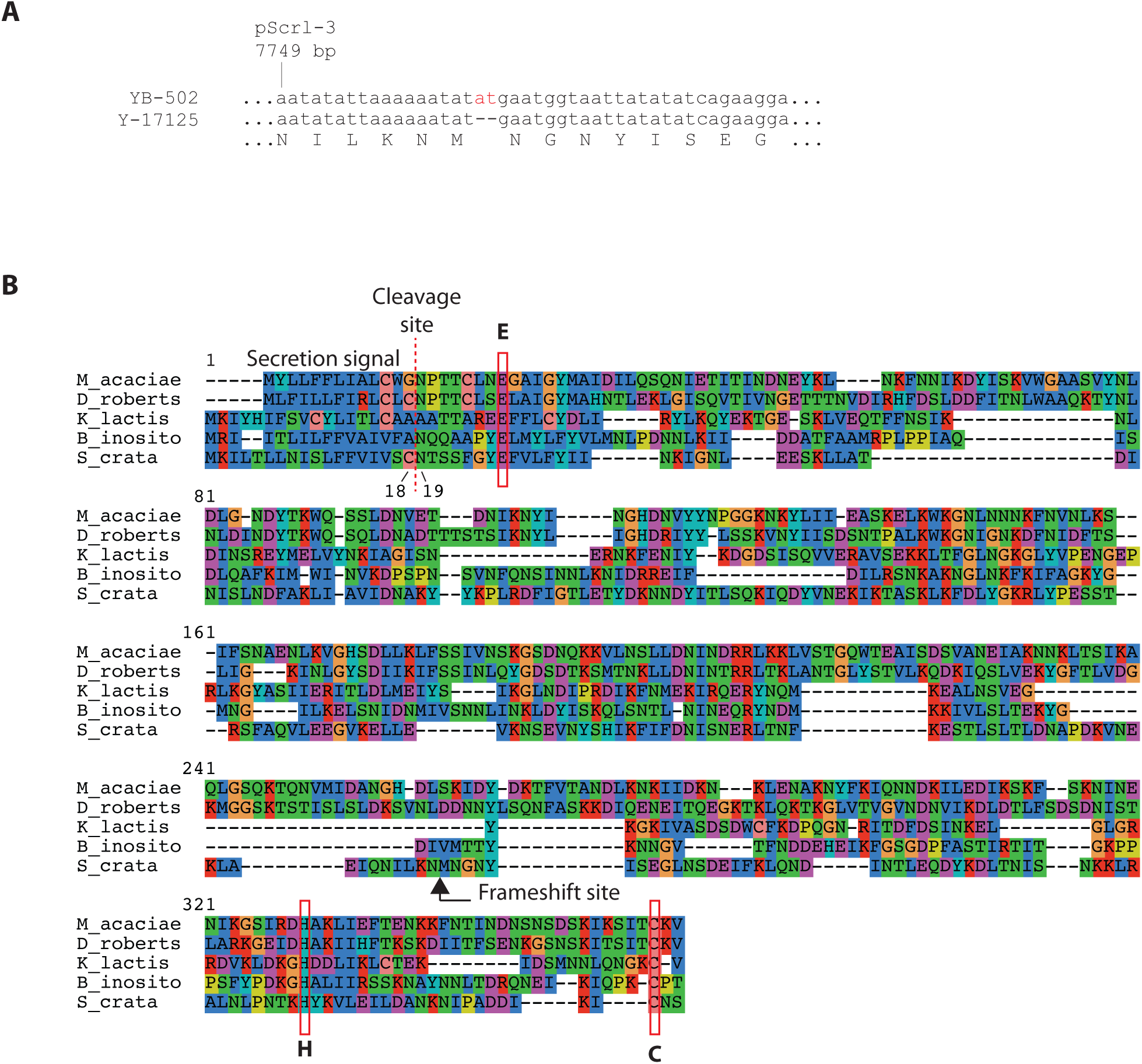
Reconstruction of the *S. crataegensis* γ-toxin sequence. (A) DNA sequence alignment of the frameshifted region of the *S. crataegensis* toxin gene from strain NRRL YB-502 (upper), and part of the toxin gene fragment in strain NRRL Y-17125 (lower) that spans the frameshift site. The inferred amino acid sequence of the region is shown. The incomplete gene in Y-17125 extends from 5 codons upstream of the frameshift site to beyond the stop codon, and an identical incomplete gene is present in strain Y-5902. (B) Amino acid sequence alignment of the reconstructed *S. crataegensis* toxin and the four known toxin sequences (PaT of *Millerozyma acaciae*, DrT of *Debaryomyces robertsiae*, zymocin γ-subunit of *Kluyveromyces lactis*, and PiT of *Babjeviella inositovora*). The predicted secretion signal sequence cleavage site and the three key residues (E, H, C) mentioned in the text are highlighted, as is the site of the corrected frameshift in *S. crataegensis*. Sequences were aligned using MUSCLE (Edgar, 2004) as implemented in Seaview version 5.04 (Gouy et al., 2010), and background coloring indicates similar amino acids. The sequences of DrT and PiT have been corrected relative to their original publications, based on resequencing of these genes in our laboratory.

### Heterologous expression in *S. cerevisiae* and *K. marxianus*

To investigate the functionality of the putative *S. crataegensis* γ-toxin, we expressed it in both *Saccharomyces cerevisiae* and *Kluyveromyces marxianus.* Both of these species use the universal genetic code and translate CUG as leucine. However, they differ in the sets of leucine tRNA genes that they contain. The ancestor of all budding yeasts used two different tRNA isoacceptors to read CUG and CUA leucine codons, without wobbling in either case: tRNA-Leu(CAG) read CUG codons, and tRNA-Leu(UAG) read CUA codons. Both of these tRNAs are still present in *K. marxianus*. In contrast, *S. cerevisiae* has no tRNA-Leu(CAG), and instead it uses tRNA-Leu(UAG) to read both CUG (by wobbling) and its cognate codon CUA (Kollmar and Mühlhausen, 2017). Thus, *K. marxianus* contains the tRNA that is the putative target of the toxin that led to the genetic code changes in budding yeasts, whereas *S. cerevisiae* does not.

We made genetic constructs that express the putative *S. crataegensis* γ-toxin gene, without its signal peptide, from an inducible promoter. This method is based on similar experiments previously used to study zymocin and PaT (Lu et al., 2005; Jablonowski et al., 2006; Chakravarty et al., 2014). The expressed toxin is retained intracellularly and has the effect of killing the host cell because immunity proteins are not present. We reasoned that if the *S. crataegensis* gene codes for a toxin that targets only tRNA-Leu(CAG) we would expect to see a growth defect phenotype only in *K. marxianus*, whereas if it targets another tRNA we would expect to see a phenotype in both *S. cerevisiae* and *K. marxianus*. However, we did not see a phenotype in either species.

For expression of the putative *S. crataegensis* γ-toxin in *S. cerevisiae*, we used the system developed by Ottoz et al. (2014), in which a synthetic transcription factor and promoter induce expression upon exposure to the human hormone β-estradiol which does not occur naturally in yeasts. The γ-toxin gene, beginning at amino acid position 19 (the predicted N-terminus of the protein after cleavage of the secretion signal; Fig. 2B), was inserted into the *S. cerevisiae* genome under the control of a β- estradiol inducible promoter (Fig. 4A). Engineered strains were streaked onto YPD plates, and onto YPD plates containing the hormone. No reduction of growth of strains expressing the *S. crataegensis* candidate toxin was observed in the presence of β-estradiol, whereas growth of control strains expressing *M. acaciae* PaT was strongly reduced in the same conditions (Fig. 4B).

**Figure 4.**
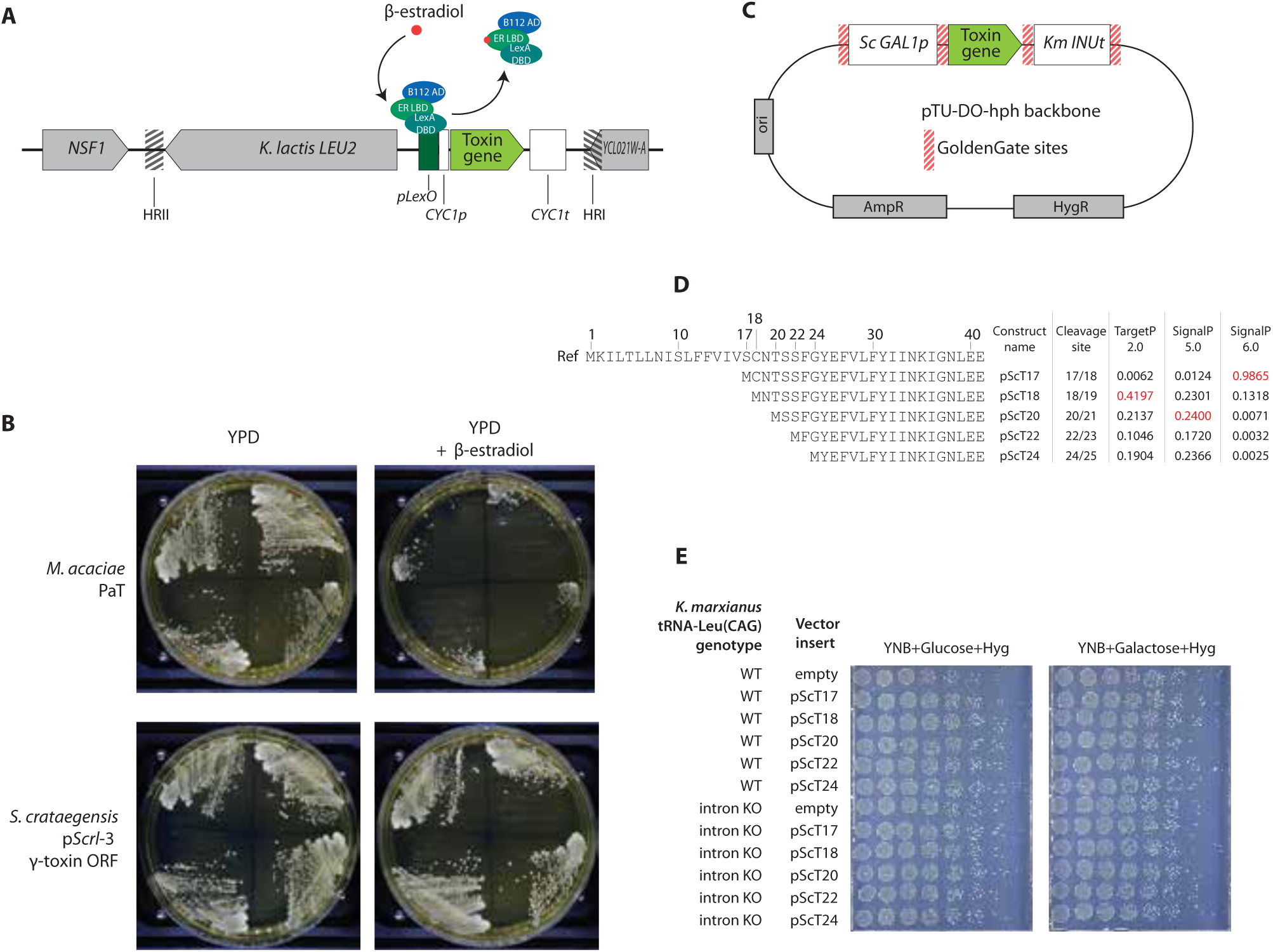
Heterologous expression of the resurrected *S. crataegensis* γ-toxin has no phenotype in *Saccharomyces cerevisiae* and *Kluyveromyces marxianus*. (A) The toxin expression construct used in *S. cerevisiae* is an integrating plasmid that complements a *leu2* mutation. It contains a synthetic promoter (*pLexO-CYC1p*) that is bound by the synthetic transcription factor LexA-ER-AD, which induces transcription of the toxin when β-estradiol is present in the media. HRI and HRII indicate sites of homologous recombination into the genome. **(B)** Induction of candidate toxins in *S. cerevisiae* by β-estradiol. The known toxin of *M. acaciae* (PaT) was used as a positive control (upper row), and the candidate *S. crataegensis* toxin was tested (lower row). Each agar plate contains quadrants with streaks of four independent clones carrying constructs as depicted in A, streaked on YPD agar (left column) or on YPD plus 2 μM β-estradiol (right column). Inhibition of yeast growth is apparent only in the *M. acaciae* clones on β-estradiol. (**C)** Map of the replicating plasmid vector constructed using the Yeast Tool Kit (YTK) for *K. marxianus*, to express the *S. crataegensis* toxin candidate under control of the *pGAL* promoter. **(D)** Amino acid sequences of N-termini of reconstructed *S. crataegensis* γ-subunit candidates expressed in *K. marxianus*. Amino acid positions are numbered from the start codon. “Ref” shows the unprocessed precursor protein. Five different possible start site variants of the mature protein are indicated, based on computational predictions of the secretion signal cleavage site by TargetP (version 5.0) and SignalP (versions 5.0 and 6.0), with site probabilities as shown on the right. Numbers in red highlight the cleavage site most favored by each computation tool. Constructs expressing each of the five variants shown were synthesized, putting a start codon immediately upstream of the predicted mature N-terminus, and assayed. **(E)** Spot assay of toxin gene variants in *K. marxianus.* Left panel, growth on glucose (no induction); right panel, growth on galactose (induction of toxin expression). Spots are serial dilutions onto agar plates containing the indicated media and hygromycin (150 μg/ml). Strain genotypes are indicated on the left. WT indicates the standard host *K. marxianus* NBRC1777 *dnl4*-1, and ‘intron KO’ indicates a derivative of this strain lacking the intron in tRNA-Leu(CAG).

For expression in *K. marxianus*, we used the GoldenGate expression chassis (Rajkumar et al., 2019), with an inducible *S. cerevisiae GAL1* promoter and *K. marxianus INU1* terminator as regulatory elements (Fig. 4C). Because the *K. marxianus* tRNA-Leu(CAG) gene contains a large intron with a complex secondary structure which might affect the interaction between the tRNA transcript and a toxin (Krassowski et al., 2018), we constructed an expression host strain in which this intron was deleted (by CRISPR/Cas9) as well as using the standard *K. marxianus* host strain NBRC1777 *dnl4*-1. As well as testing a construct that began at amino acid 19 of the toxin as used in *S. cerevisiae*, we also tested constructs that began at four other positions that were predicted as possible alternative cleavage sites by either the TargetP or SignalP programs (Emanuelsson et al., 2007) (Fig. 4D). Toxin activity was assayed by serial dilution onto plates containing galactose (to induce expression of the toxin) or glucose as a control. However, none of the constructs caused a significant reduction of growth of *K. marxianus* (Fig. 4E), regardless of the toxin N-terminus used or the presence/absence of the tRNA-Leu(CAG) intron. We conclude that the resurrected *S. crataegensis* toxin is inactive against both *S. cerevisiae* and *K. marxianus*. Since the gene was not fully intact in any of the *S. crataegensis* strains we examined, it is possible that the gene sequence we used contains other inactivating mutations such as missense mutations, in addition to the frameshift we corrected.

### Diversity of plasmid DNA polymerase genes

The newly sequenced plasmids of this study greatly expand our knowledge of cytosolic linear DNA plasmid diversity, and permit greater investigation of the evolutionary history of these types of elements within the budding yeasts. We used the DNA polymerase (DNAP) genes, which are present in many plasmids (Fig. 1) and relatively slowly evolving, as a phylogenetic marker. We constructed a phylogenetic tree from an alignment of DNAP proteins from *Saccharomycopsis* plasmids and from all other currently sequenced yeast cytosolic linear DNA plasmids (Fig. 5). IQ-Tree (Nguyen et al., 2015) was used to find the best model and build the resulting maximum likelihood tree, measuring branch support by ultrafast bootstrapping and SH-arlt tests. Several branches are not highly supported, but the overall topology of the tree is clear and several conclusions can be drawn from it.

**Figure 5.**
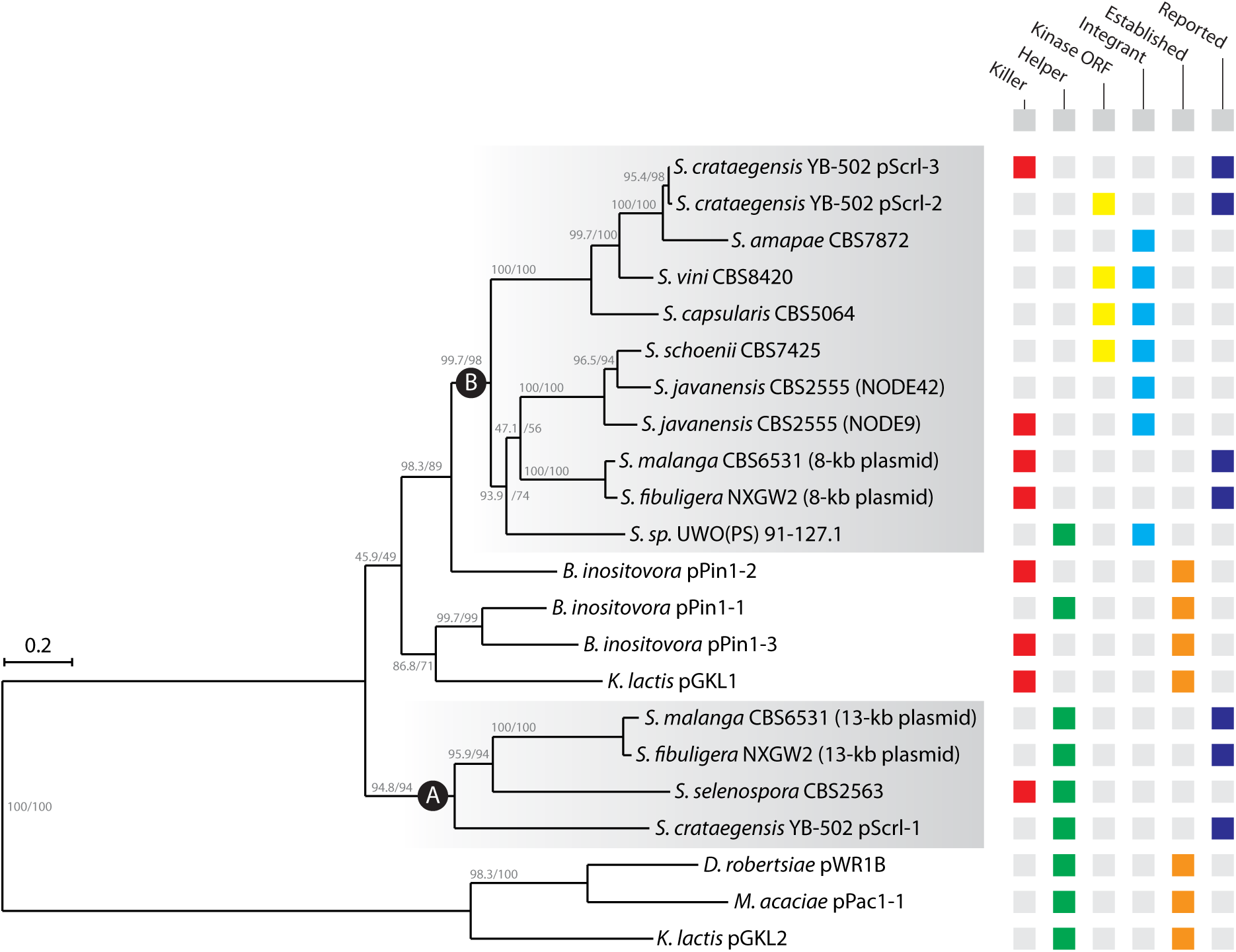
Maximum likelihood phylogeny of DNA polymerase proteins encoded by cytosolic linear DNA plasmids or plasmid-like nuclear integrations, from *Saccharomycopsis* and other species. The grid on the right summarizes the context in which the DNA polymerase gene was found. Killer: the contig contains genes or pseudogenes for predicted toxin α/β (chitin-binding) or γ (ribonuclease) subunits. Helper: large plasmid containing housekeeping genes such as those in pGKL2. Kinase ORF: gene with kinase (phosphotransferase) conserved domain is present. Integrant: the plasmid element is likely integrated into the nuclear genome. Established: the plasmid is well known and previously investigated. Reported: plasmids that were newly sequenced in this study but were detected by gel electrophoresis in previous studies. The tree was constructed using IQ-Tree. Two types of branch support values (bootstrap/SH-alrt) are shown for each internal branch.

First, the DNAPs of the helper plasmids of the established killer systems in *K. lactis* (pGKL2), *D. robertsiae* (pWR1B), and *M. acaciae* (pPac1-1) form an outgroup to all the other DNAPs. Second, the DNAPs of most of the helper plasmids in *Saccharomycopsis* species form a clade (labelled A in Fig. 5) that is distinct from the outgroup helper plasmids and more closely related to a clade of DNAPs from killer plasmids. Within the latter clade, there is a monophyletic group (labelled B) of DNAPs that are exclusively from *Saccharomycopsis* species. For the plasmid systems we characterized in *S. crataegensis*, *S. malanga* and *S. fibuligera*, the helper plasmid DNAP is in clade A, and the other plasmid DNAP(s) are in clade B. Clade B is also interesting because it contains all the instances of a kinase (ORF_K) homolog, whose function is unknown and which is not present in any of the four well-characterized toxin plasmid systems. Third, the helper plasmid DNAPs from *Saccharomycopsis* UWOPS 91-127.1 and *Babjeviella inositovora* pPin1-1 fall outside the main clade of helper plasmid DNAPs (clade A) and are instead closer to killer plasmid DNAPs from the same genus (in clade B), which is suggestive of horizontal gene transfer or gene conversion between the DNAPs of different plasmids. Fourth, the large 21.8-kb plasmid of *S. selenospora*, which resembles a fusion between a helper plasmid and a killer plasmid (Fig. 1D), retains only a helper plasmid-type DNAP (clade A). Overall, the tree indicates a previously hidden diversity of plasmid systems contained solely within *Saccharomycopsis*, that have been evolving independently of the better-known plasmids in other yeasts.

### Conclusions

We identified and sequenced the cryptic linear DNA plasmids of several *Saccharomycopsis* species and found that they have a close yet distinct relationship with other extant plasmids of this type. Only one plasmid contains a potential γ-toxin gene, which we could not show is a functional toxin. It remains possible that its amino acid sequence was inferred incorrectly, or that the target species specificity may lie outside the species we tested. The plasmids of *Saccharomycopsis* are diverse, have been evolving for some time, and show some evidence of horizontal transmission between species. The presence of integrated plasmid-derived sequences in the nuclear genomes of three species (*S. capsularis*, *S. vini*, *S. javanensis*) in which free-living plasmids have not yet been detected suggests that the genus contains (or contained) yet more undiscovered diversity of plasmids. The presence of genes with kinase (aminoglycoside 3’-phosphotransferase) domains is the first instance of such genes being present on plasmids of this type, and their significance is unknown.

Furthermore, the heterogeneity that we found within the single species *S. crataegensis* (we only found a complete set of full-length plasmids in one of the nine strains we sequenced), and similar heterogeneity in other species (Fukuhara, 1995; Polomska et al., 2021), emphasizes the need to sequence broad panels of environmental isolates to fully uncover the plasmid repertoire of any given species.

## Acknowledgements

This work was supported by the European Research Council (789341). We thank Jürgen Wendland for sharing data from *Saccharomycopsis* UWOPS 91-127.1.

## Materials and Methods

### Genome sequencing and assembly

Yeast strains sequenced and/or analyzed in this study are listed in Table S1. Our choice of *Saccharomycopsis* strains for sequencing was primarily guided by previous reports of presence of dsDNA linear plasmids (Shepherd et al., 1987; Fukuhara, 1995). Strains were obtained from the USDA ARS culture collection (NRRL), the Westerdijk Institute for Fungal Diversity (CBS Collection, The Netherlands), and the Portuguese Yeast Culture Collection (PYCC). We were unable to obtain two *S. crataegensis* strains (NRRL Y-5903 and NRRL Y-5910) that were previously studied by Bolen et al. (1992).

We carried out whole-genome sequencing and did not try to isolate plasmid DNA. Cultures of all strains were grown in YPD at 30°C overnight. Cells were harvested by centrifugation and cell pellets were resuspended in 200 µl extraction buffer (2% Triton X100, 100 mM NaCl, 10 mM Tris pH 7.4, 1 mM EDTA, 1% SDS) in a 1.5 ml screw-cap tube. Approximately 0.3 g acid-washed glass beads (425- 600 μm) were added with 200 μl phenol/chloroform/isoamyl alcohol (25:24:1). The mixture was agitated on a 600 MiniG bead beater (Spex SamplePrep) at 1,500 rpm, 4-6 times for 30 s each, and centrifuged at 15,000 rcf for 10 min. The top aqueous layer was transferred to a new 1.5 ml screw-cap tube, 200 μl TE buffer was added, and 200 μl of the phenol/chloroform/isoamyl alcohol mixture was added. This was agitated as before on the 600 MiniG bead beater, centrifuged at 14,000 rpm for 10 min. The top aqueous layer was transferred to a new microfuge tube, after which 80 μl 7.5 M ammonium acetate and 1 ml 100% isopropyl alcohol were added to precipitate the DNA. DNA was pelleted by centrifugation at 14,000 rpm, washed using 70% ethanol and dried in a SpeedVac (Eppendorf Concentrator 5301 at 45°C for 2 min pulses until dry). Pellets were resuspended in 400 µl TE buffer with 1 µl RNase A (10 mg/ml) and incubated overnight at 37°C. DNA was re-precipitated and washed once more as above and re-suspended in 150 µl water. DNA quality and concentration was assessed by gel electrophoresis, Nanodrop, and Qubit measurement. Genomic DNA was sequenced by the UCD Conway Institute Genomics Core Facility (Ireland) using Illumina NextSeq 500 and 550 instruments (150 bp paired ends, 1.5 Gb raw data per sample).

Genome sequences were assembled using SPAdes version 3.11 (Bankevich et al., 2012). Reference sequences for the three plasmids in *S. crataegensis* NRRL YB-502 were assembled by using SPAdes with the custom parameters -k 127 -c 100, followed by manual joining of overlapping contigs. All other genomes sequenced in this study, except *S. selenospora* CBS 2563, were assembled using SPAdes with default parameters, which resulted in the sequences of the p*Scr*l plasmids being fragmented into many small contigs in our assemblies of *S. crataegensis* strains Y-5902, Y-5904, Y- 17125 and PYCC 8280. The plasmid structures in these strains were inferred by alignment of contigs to the NRRL YB-502 reference. The 21.8 kb plasmid in *S. selenospora* CBS 2563 was also assembled using the custom parameters -k 127 -c 100. In addition to assembling our own Illumina data, we also made *de novo* assemblies of 93 *S. fibuligera* strains for which unassembled Illumina reads were deposited in the SRA database by Wang et al. (2021). Among these, we identified both the 13-kb and 8-kb plasmid in strains NXGW2 (SRR15179763) and MTDQ3 (SRR15179688), and only the 8-kb plasmid in strain CGMCC2.0968 (SRR15179711).

New genome sequences reported in this study have been deposited in the NCBI/ENA/DDBJ databases under BioProject number PRJNA977123.

### Heterologous expression of *S. crataegensis* gamma subunit in *S. cerevisiae*

The inferred intact toxin γ-subunit ORF of *S. crataegensis* p*Scr*l-3 was synthesized as a synthetic DNA fragment by Twist Biosciences and cloned into the β-estradiol inducible plasmid pRG634 (Ottoz et al., 2014; Gnugge and Symington, 2020). The plasmid was integrated into *S. cerevisiae* and four independent transformants were taken forward for toxin screening. Strains were streaked to YPD or YPD +2 μM β-estradiol and compared visually two days later. A parallel experiment was performed with the PaT toxin of *M. acaciae* as a positive control.

### *K. marxianus* CRISPR gene editing and heterologous expression

Guide RNA sequences were designed targeting the intron of tRNA-Leu(CAG) in *K. marxianus*. Two guides (5’-TACAAAAGTTCAGCTTTGCT-3’ and 5’-AAAACCAAGGTGCAGCTCCA-3’) were cloned into the CRISPR plasmid (Rajkumar et al., 2019; Rajkumar and Morrissey, 2020) and transformed into *K. marxianus* ASR.005 (Rajkumar et al., 2019), along with a dsDNA repair template with 75 bp homology to either side of the intron. The repair template was generated by primer extension. Transformants were screened by PCR across the tRNA-Leu(CAG) locus and confirmed by a shift in band size (approximately 200 bp), and 2 independent transformants from each guide plasmid were cultured in YPD broth to cure the strains. After two days, a 1e-06 dilution of this culture was plated to YPD agar to isolate cured single colonies.

A synthetic gene for the reconstructed *S. crataegensis* toxin ORF, codon optimized for *K. lactis* (a close relative of *K. marxianus*) and beginning at codon 19, was synthesized by Integrated DNA Technologies. We used PCR oligonucleotides to generate the other four start site variants (Fig. 4D). All five constructs were assembled by the *K. marxianus* GoldenGate system into backbone vector pTU-DO-hph (Rajkumar et al., 2019; Rajkumar and Morrissey, 2020), with the *S. cerevisiae GAL1* promoter and *K. marxianus INU1* terminator. These plasmids, and empty vector controls, were transformed into two separate *K. marxianus* strains: one with an intact tRNA-Leu(CAG), and one in which the intron of tRNA-Leu(CAG) had been deleted.

For spot assays, overnight cultures (5 ml) of transformants were grown with hygromycin selection (150 μg/ml), diluted to the same OD600, and then manually pinned on agar plates containing YNB + glucose + hygromycin or YNB + galactose + hygromycin. Plates were incubated at 30°C for 2 days, then imaged.

### DNA polymerase phylogeny

DNA polymerase genes from known and newly discovered plasmids were collected. Only those DNA polymerase genes that were fully or mostly intact were used for alignment and subsequent tree construction. The sequences were aligned with MUSCLE (default) and IQ-tree v1.6.12 (Nguyen et al., 2015) was used to find the best model (WAG+F+I+G4) and construct a tree.

---

**Supplementary** Figure 1. Phosphotransferase conserved domains in ORF_K. (A) Structure of conserved domains (APH_ChoK_like in orange; CotS in navy) within ORF_K of p*Scr*l-2. Numbers refer to amino acid positions. The dashed red line indicates the position of an internal in-frame stop codon. (B) Alignments of conserved domain sections with their hit in the NCBI Conserved Domains Database (CDD). Conserved residues are highlighted in gry. BLAST E-values are shown at the top-right of each alignment.

