## Supplementary Information for "Cytosolic linear DNA plasmids in *Saccharomycopsis* species"

**A**

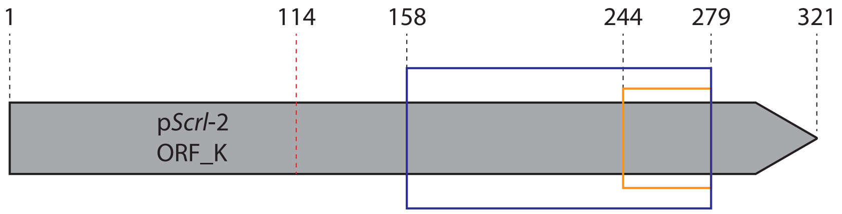


**B**

APH_ChoK_like
Aminoglycoside 3'-phosphotransferase 2.12e-06

```
     ORF_K 244 VLSHGDLHPENIIVKNDLSVY-FIDWENMGLLPDYWD 279
Cdd:cd05120 112 VLTHGDLHPGNILVKPDGKLSgIIDWEFAGYGPPAFD 148
```

CotS
Thiamine kinase 1.01e-05

```
     ORF_K 158 INGETLSKICSSLNKEEIVIVSNNIIKEIIKLRNIK----------FDKIVEI 200
Cdd:COG0510  71 ENGRTLTPEDMNLKK----------IAHILKKLHNSVPLLHQLPRSGSSFIEP 113

           201 KDYIYGELKFKNEKELNKNVCEFVKNKIEKDIIIELLPKNSKFVLSHGDLHP 252
           114 KDYL--ELLWQQNSRAYRDNHLLRKKLKELRRALEEVPKDDL-VPCHNDLNP 162

           253 ENIIVKNDLSVYFIDWENMGLLPDYWD 279
           163 GNLLLTDKGGLFLIDWEYAGLNDPAFD 189
```

Supplementary Figure 1

**Table S1.** *Saccharomycopsis* strains sequenced or analysed in this study.

| Species | Strain | Plasmid elements found in genome sequence | Plasmids reported by gel electrophoresis | Accession numbers and reference for genome sequence |
|---|---|---|---|---|
| *S. crataegensis* | NRRL YB-502 | p*Scr*l-1 (13.3 kb)<br>p*Scr*l-2 (7.2 kb)<br>p*Scr*l-3 (9.9 kb) | Not examined | JASTUR000000000 (this study) |
| | NRRL Y-5902<br>(type strain) [a] | p*Scr*l-1<br>p*Scr*l-2<br>p*Scr*l-3 [b] (Δ 4 kb) | p*Scr*l-1<br>p*Scr*l-2<br>p*Scr*l-3<br>(Bolen, et al., 1992) | JASTUU000000000 (this study) |
| | NRRL Y-5904 | p*Scr*l-1<br>p*Scr*l-2<br>p*Scr*l-3 [b] (Δ 6 kb) | p*Scr*l-1<br>p*Scr*l-2<br>(Bolen, et al., 1992) | JAUJEI000000000 (this study) |
| | NRRL Y-17125 | p*Scr*l-1<br>p*Scr*l-2<br>p*Scr*l-3 [b] (Δ 4 kb) | Not examined | JASTUT000000000 (this study) |
| | NRRL YB-192 | None | None (Bolen, et al., 1992) | JASTUS000000000 (this study) |
| | PYCC 8280 | p*Scr*l-1 only | Not examined | JAUJEJ000000000 (this study) |
| | CBS 7190 | None | None (Fukuhara, unpublished data on CBS website) | JAUJEK000000000 (this study) |
| | CBS 6447<br>(type strain) [a] | None | Yes (synonym of NRRL Y-5902) | JASTUW000000000 (Ó Cinnéide, et al., 2023) |
| | CBS 6448 | None | Yes (synonym of NRRL Y-5904) | JASTUV000000000 (Ó Cinnéide, et al., 2023) |

| | | | | |
|---|---|---|---|---|
| *S. fibuligera* | CBS 7434 | 13.2 kb, 8 kb | Yes (Fukuhara, 1995) | JASTUO000000000 (this study) |
| | NXGW2 | 13.2 kb, 8 kb | No | (Wang, et al., 2021), reads assembled in this study (SRA accession SRR15179763) |
| *S. malanga* | CBS 6531 | 13.1 kb, 8 kb | Yes (Fukuhara, 1995) | JASVEE000000000 (this study) |
| *S. selenospora* | CBS 2563 | 21.8 kb | Yes (Fukuhara, unpublished data on CBS website) | JASTUF000000000 (this study) |
| *Saccharomycopsis*, unnamed species. | UWO(PS) 91-127.1 | 14.2 kb (integrated in nuclear genome) | No | JNNM01000000 (J. Wendland & A. Hesselbart, unpublished) |
| *S. capsularis* | CBS 5064 | 4.4 kb (fragment integrated in nuclear genome) | No | JASTUZ000000000 (Ó Cinnéide, et al., 2023) |
| *S. vini* | CBS 8420 | 4.3 kb (fragment integrated in nuclear genome) | No | JASTUA000000000 (Ó Cinnéide, et al., 2023) |
| *S. amapae* | CBS 7872 | 3.3 kb (fragment integrated in nuclear genome) | No | JASTVC000000000 (Ó Cinnéide, et al., 2023) |
| *S. javanensis* | CBS 2555 | 4.3 and 5.7 kb (fragments integrated in nuclear genome) | No | JASTUL000000000 (Ó Cinnéide, et al., 2023) |
| *S. schoenii* | CBS 7425 | 4.5 kb (fragment integrated in nuclear genome) | No | JNFU01000000 (Junker, et al., 2019) |

[a] *S. crataegensis* strains NRRL Y-5902 and CBS 6447 are duplicate stocks of the type strain of *S. crataegensis*, held in the US Department of Agriculture (NRRL) and Westerdijk Institute (CBS) culture collections respectively. They are expected to be identical but CBS 6447 appears to have lost the plasmids.

[b] The putative killer plasmid p*Scr*l-3 is shorter in these three strains than in our reference strain NRRL YB-502. Strains Y-5902 and Y-17125 lack a 4-kb region (positions 3754 to 7747 in the YB-502 sequence) spanning most of the α/β subunit gene and part of the γ subunit gene, and strain Y-5904 lacks the 6 kb from position 3754 in the α/β subunit gene to the right-hand end of the plasmid, including the whole γ-toxin gene.

Bolen, P. L., Kurtzman, C. P., Ligon, J. M., Mannarelli, B. M. and Bothast, R. J. (1992). Physical and genetic characterization of linear DNA plasmids from the heterothallic yeast Saccharomycopsis crataegensis. *Antonie Van Leeuwenhoek* **61,** 195-205.
Fukuhara, H. (1995). Linear DNA plasmids of yeasts. *FEMS Microbiol Lett* **131,** 1-9.
Junker, K., Chailyan, A., Hesselbart, A., Forster, J. and Wendland, J. (2019). Multi-omics characterization of the necrotrophic mycoparasite Saccharomycopsis schoenii. *PLoS Pathog* **15,** e1007692.
Ó Cinnéide, E., Scaife, C., Dillon, E. and Wolfe, K. H. (2023). A genetic code change in progress: tRNA-Leu(CAG) is conserved in most *Saccharomycopsis* yeast species but is non-essential and does not compete with tRNA-Ser(CAG) in translation. *BioRxiv*.
Wang, J. W., Han, P. J., Han, D. Y., Zhou, S., Li, K., He, P. Y., Zhen, P., Yu, H. X., Liang, Z. R., Wang, X. W. and Bai, F. Y. (2021). Genetic diversity and population structure of the amylolytic yeast *Saccharomycopsis fibuligera* associated with Baijiu fermentation in China. *J Microbiol* **59,** 753-762.
